# Ant cuticular hydrocarbons are heritable and associated with variation in colony productivity

**DOI:** 10.1101/819870

**Authors:** Justin Walsh, Luigi Pontieri, Patrizia d’Ettorre, Timothy A. Linksvayer

## Abstract

In social insects, cuticular hydrocarbons function in nestmate recognition and also provide a waxy barrier against desiccation, but basic evolutionary genetic features, including the heritability of hydrocarbon profiles and how they are shaped by natural selection are largely unknown. We used a new pharaoh ant (*Monomorium pharaonis*) laboratory mapping population to estimate the heritability of individual cuticular hydrocarbons, genetic correlations between hydrocarbons, and fitness consequences of phenotypic variation in the hydrocarbons. Individual hydrocarbons had low to moderate estimated heritability, indicating that some compounds provide more information about genetic relatedness and can also better respond to natural selection. Strong genetic correlations between compounds are likely to constrain independent evolutionary trajectories, which is expected given that many hydrocarbons share biosynthetic pathways. Variation in cuticular hydrocarbons was associated with variation in colony productivity, with some hydrocarbons experiencing strong directional selection. Altogether, our study builds on our knowledge of the genetic architecture of the social insect hydrocarbon profile and demonstrates that hydrocarbon variation is shaped by natural selection.

## Introduction

Cuticular hydrocarbons play several critical roles in social insect societies. As in solitary insects, social insect cuticular hydrocarbons provide a waxy barrier that prevents desiccation, and they are also chemical signals used in kin and species recognition during mate choice. Furthermore, they are perhaps best studied as being the main chemical signals used in nestmate recognition (Bonavita-Cougourdan et al. 1987; Lahav et al. 1999; Lenoir et al. 1999; Wagner et al. 2000; Greene & Gordon 2007; Martin & Drijfhout 2009a; van Zweden & d’Ettorre 2010). In many social insect species, they also encode information about the reproductive status, dominance status, and current task of individuals within colonies (Edwards & Chambers 1984; Wagner et al. 1998, 2001; Green & Gordon 2003; Martin & Drijfhout 2009a; Holman et al. 2010; Liebig 2010; Abril et al. 2018).

Despite the central role of cuticular hydrocarbons in insect societies, we still know very little about fundamental genetic and evolutionary features shaping social insect hydrocarbon profiles, including the relative contribution of genetic and environmental factors to phenotypic variation in hydrocarbon profiles within and between colonies (Menzel et al. 2017), and how natural selection acts on this variation. Numerous studies have demonstrated that the social insect hydrocarbon profile is influenced by genotype by tracking familial lines (van Zweden et al. 2009; Nehring et al. 2011; Martin et al. 2012, Martin et al. 2013), using cross-fostering designs (van Zweden et al. 2010), or demonstrating an association between hydrocarbon diversity and within-colony genetic variation (Dronnet et al. 2006; Menzel et al. 2016). However, very few studies have examined the underlying genetic architecture of social insect hydrocarbons within a formal quantitative genetic framework (Boomsma et al. 2003). Traditional quantitative genetic crossing and pedigree-based mapping populations provide a powerful means to elucidate the contribution of genetic and environmental factors to variation in hydrocarbon profiles (e.g. Thomas & Simmons 2008; Sharma et al. 2012; Dembeck et al 2015, Berson et al. 2019).

The genetic architecture of social insect cuticular hydrocarbons is expected to be more complex than solitary insect cuticular hydrocarbons because the hydrocarbon profile of each individual can be made up of compounds synthesized directly by the individual itself, as well as compounds synthesized by nestmates and socially transferred to the individual (van Zweden & d’Ettorre 2010; van Zweden et al. 2009, 2010; Linksvayer 2015; Leonhardt et al. 2016). More generally, in social organisms such as social insects, an individual’s own traits can be influenced directly by its own genotype (i.e. direct genetic effects) but also indirectly via the genotype of social partners (i.e. indirect genetic effects) (Linksvayer 2006, 2015). Indeed, hydrocarbons are known to be transferred among colonies members via allogrooming and the exchange of regurgitated liquids (i.e. trophallaxis) (van Zweden & d’Ettorre 2010; van Zweden et al. 2009, 2010; Leonhardt et al. 2016), creating a collective colony odor that allows workers to distinguish between nestmates and non-nestmates (Crozier & Dix 1979).

To fully understand the hydrocarbon profile’s potential evolutionary response to natural selection, in addition to understanding quantitative genetic parameters such as heritability and genetic correlations, we must also understand the fitness consequences of phenotypic variation in the hydrocarbon profile. Knowledge of the fitness consequences of trait variation is necessary to characterize the type (e.g., directional, stabilizing, or disruptive) and strength of natural selection acting on a trait (Lande & Arnold 1983; Arnold & Wade 1984; Janzen & Stern 1998; Morrissey & Sakrejda 2013). Variation in the social insect hydrocarbon profile may affect individual survival and colony productivity by affecting desiccation resistance (Gordon 2013; Buellesbach et al. 2018; Sprenger et al. 2018; Friedman et al. 2019) or by influencing chemical communication among nestmates and the collective behavior of the colony (Lahav et al. 2001; Green & Gordon 2003; Martin & Drijfhout 2009a; Liebig 2010; Buellesbach et al. 2018).

Hydrocarbon structural classes (i.e. alkenes, linear alkanes, monomethyl alkanes, and dimethyl alkanes) have distinct functional properties that are likely to influence the roles they play in insect societies and how they are shaped by natural selection (Martin & Drijfhout 2009b; Menzel et al. 2017. Linear alkanes provide the best desiccation resistance because these molecules tightly aggregate (Gibbs 1995, 1998; Gibbs & Rajpurohit 2010). On the other hand, alkenes and monomethyl and dimethyl alkanes are expected to play a larger role in chemical communication because they can be distinguished by the position of their double bond or of the methyl group(s), while linear alkanes can only be distinguished based on their chain length (Gibbs & Pomonis 1995; Akino et al. 2004; Dani et al. 2005; Martin et al. 2008). This increased complexity allows alkenes and monomethyl and dimethyl alkanes to encode more information and likely play a stronger role in chemical communication when compared to linear alkanes. There is evidence that linear alkanes are less heritable and not transferred between workers as much as monomethyl and dimethyl alkanes, suggesting that linear alkanes are less informative for nestmate recognition (van Zweden et al. 2010). Furthermore, hydrocarbons of different functional classes and length share distinct biosynthetic pathways and hence may show distinct patterns of heritability and also genetic correlations within and between different types of hydrocarbons.

Here, we use a genetically highly variable laboratory population of pharaoh ant (*Monomorium pharaonis*) colonies that we created by systematically intercrossing eight initial parental lineages for nine generations (Walsh et al. 2019). Such a genetically heterogeneous mapping population has proven powerful to elucidate the genetic architecture of a range of traits in rats, mice, and fruit flies (e.g., Hansen & Spuhler 1984; Mott et al. 2000; Valdar et al. 2006; King et al. 2012; de Koning & McIntyre 2017). We first extract hydrocarbons from three groups of 15 workers (45 workers total) from three replicate sub-colonies of 48 distinct colony genotypes of known pedigree. Next, we use the pedigree information of the colonies to estimate the heritability of and the genetic correlations between hydrocarbons. Additionally, we use a random forest analysis to identify hydrocarbons that best discriminate between our *M. pharaonis* colony genotypes. Finally, we estimate the strength and pattern of natural selection putatively acting on hydrocarbons in the laboratory.

## Methods

### Experimental design and colony maintenance

We used 48 lab reared *M. pharaonis* colonies (hereafter “colony genotypes”) of known pedigree from our heterogeneous stock mapping population, which was derived from eight initial lab stock colonies that were systematically interbred for nine generations (Supplementary figure 1, see Walsh et al. 2019 and Pontieri et al. 2017 for details). We split each colony genotype into three equally sized replicates (hereafter “colony replicates”) that initially consisted of 4 queens, 400 ± 40 workers, 60 ± 6 eggs, 50 ± 5 first instar larvae, 20 ± 2 second instar larvae, 70 ± 7 third instar larvae, 20 ± 2 prepupae, and 60 ± 6 worker pupae (Supplementary figure 2). These colony demographics represent a typical distribution found in a relatively small *M. pharaonis* colony (Buczkowski & Bennett 2009; Warner et al. 2018).

We maintained all colony replicates on a 12:12 hour light:dark cycle and at 27 ± 1 °C and 50% relative humidity. We fed each colony replicate twice per week with an agar-based synthetic diet (Dussutour & Simpson 2008) and mealworms. Water was provided *ad libitum* via a glass tube plugged with cotton (Walsh et al. 2018). Colony replicates nested between two glass slides (5 cm x 10 cm) housed in a plastic container (18.5 cm x 10.5 cm x 10.5 cm) lined with Fluon^Ⓡ^.

### Behavioral and colony productivity data collection

We surveyed each colony replicate for five collective behaviors and two measures of colony productivity (Supplementary figure 2). First, we established five assays to collect measurements of the following collective behaviors (see Walsh et al. 2019 for assays design details): 1) foraging (defined as the number of workers that visited a food source in an hour); 2) aggression (the number of aggressive acts observed in an hour towards workers of a second species, *Monomorium dichroum*); 3) exploratory rate (a measure of how quickly workers disperse from the nest during the exploration of a novel arena, Börger & Fryxell 2012); 4) group exploration (the percent of a novel arena explored by a group of five foragers in 15 minutes) and 5) colony exploration (the percent of a novel arena explored by the entire colony in 15 minutes).

After we completed the behavioral assays, the queens from each colony replicate were removed to trigger sexual production (Edwards 1991; Warner et al. 2018). We conducted weekly surveys of the number of worker, gyne (i.e. virgin queens) and male pupae produced, until all brood matured. We used the 1) total number of sexual pupae (i.e. gyne + males) and 2) the total number of worker pupae as final measures of colony productivity.

Behavioral and colony productivity data for each colony replicate are reported in **Supplementary table 1**.

### Collection, extraction and analysis of cuticular hydrocarbon samples

Upon completion of behavioral and colony productivity surveys, from each colony replicate we collected three samples, each consisting of 15 workers, for the extraction and analysis of cuticular hydrocarbons (Supplementary figure 2). To reduce the secretion of glandular compounds (e.g., alkaloids) that workers emit in response to disturbance and to facilitate the access to different nest areas, the colony was briefly anesthetized with CO_2_ and the upper glass lid covering the nest was removed prior collection. To ensure that each sample represented the average cuticular hydrocarbon profile of the whole colony, we collected a mix of of workers of different ages and tasks (i.e. nurses and foragers) by assessing cuticle coloration and position in the nest (in *M. pharaonis* nurses, which are young and light-brown in color, tend the brood located in the central area of the nest, whereas foragers, older and dark-brown, spend more time at the nest periphery or in the foraging area outside the nest. See Mikheyev & Linksvayer 2015). We took particular care to not collect the callow workers, easily detectable due their extreme pale coloration, as their cuticle likely do not possess all the hydrocarbon compounds of a mature individual (Dahbi et al. 1998). We collected workers using clean forceps, which were rinsed in pentane after sampling from each colony replicate to avoid transfer of compounds across samples of different colonies. Each group of 15 workers was promptly placed in a clean petri dish and killed by storing in a freezer at −20 °C for 20 minutes.

To extract the cuticular hydrocarbon profile, we swiftly transferred each group of workers into a clean 2 mL glass vial (Supelco) and rinsed in 200 μl of HPLC grade (99%) pentane (Sigma-Aldrich) for 10 min. We transferred the extract in a 0.2 mL glass insert (Sigma-Aldrich), removed the ants, placed the insert in the glass vial and let the extract evaporate under a fume hood. The dried extract was then stored at −20 °C until use. Immediately prior to analysis we resuspended the dried extract into 10 μl of pentane, and an aliquot (3 μl) of the solution was analyzed.

We carried out capillary gas chromatography (GC) using an Agilent 6890N gas chromatograph equipped with a HP-5MS column (length: 30 m; ID: 0.25 mm; film thickness 0.25 μm) and coupled with a 5375 Agilent mass spectrometer with 70 eV electron impact ionization. The injector was split-splitless with helium as the carrying gas at 1 mL/min. We set the initial oven temperature to 70°C, and then increasing it from 70 °C to 280 °C at 20 °C/min^−1^ and from 280 °C to 310 °C at 2 °C/min^−1^, finishing with a hold at 310 °C for 5 min. We identified 34 chemical compounds (Figure 1a) by their retention times, fragmentation patterns and comparison with published results (Schmidt et al. 2010; van Zweden 2014). We integrated the area under each peak using MSD Chemstation (Agilent Technologies, Santa Clara, CA, U.S.A.). As they had similar retention times, some compounds co-eluted into the same peak (Peak 2: y-C_25:1_ with Peak 3: *n*-C_25_; Peak 27: x-C_31:1_ with Peak 28: y-C_31:1_; Figure 1a). We combined the areas of each co-eluting pair (Peak 2+3 and Peak 27+28, respectively), leaving 32 peaks available for statistical analysis.

**Figure 1.**
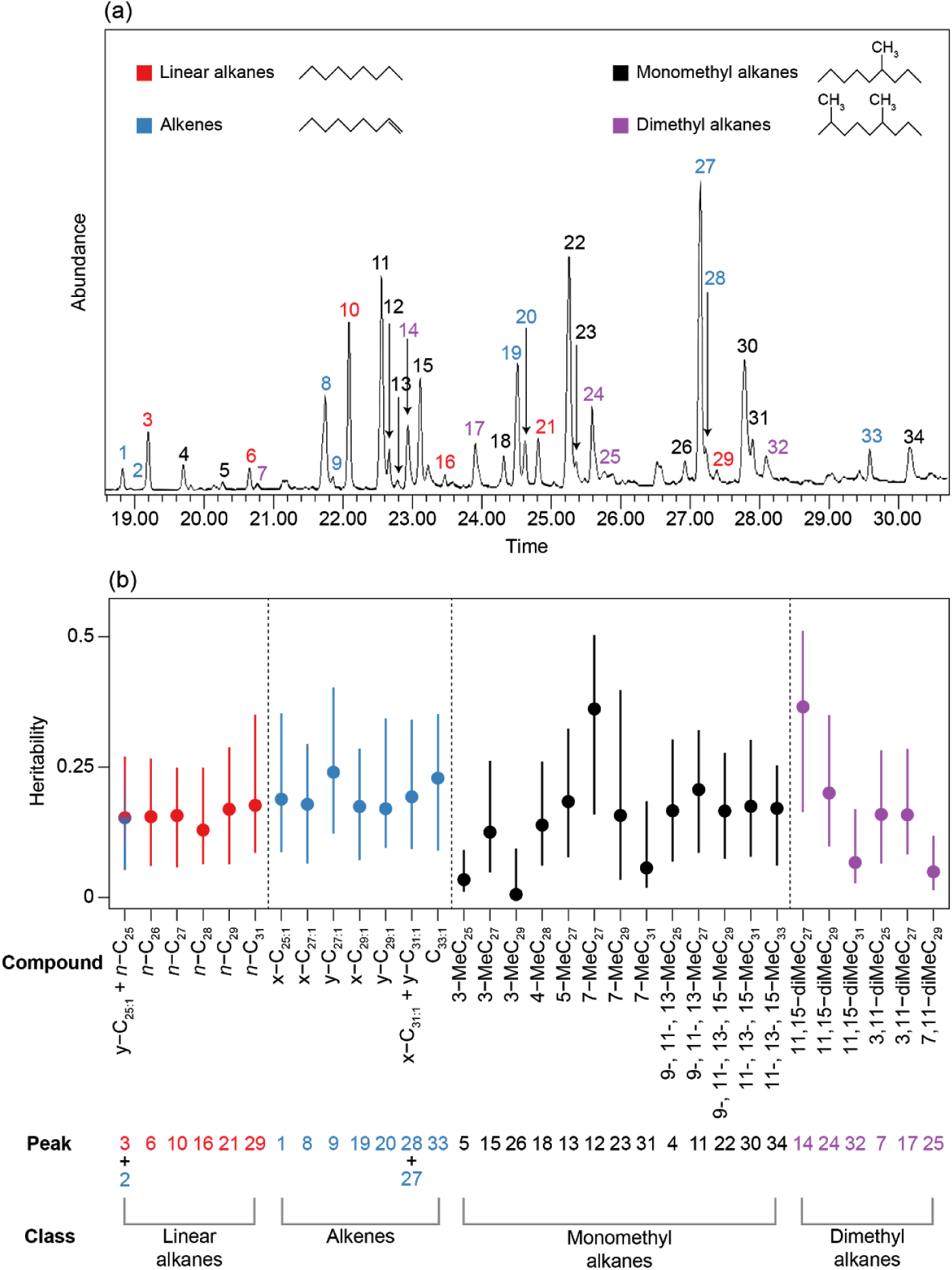
(a) GC-MS spectrum of *M. pharaonis* cuticular hydrocarbons. All 34 identified peaks are numbered and color-coded by structural class. Unidentified peaks were either contaminants or unidentifiable compounds. (b) Caterpillar plot showing heritability estimates of individual hydrocarbon compounds, with associated 95% confidence intervals, obtained from univariate animal models. Peak numbers showed in (a) are reported below the corresponding compound(s). Compounds in the plot are grouped and color-coded by structural class and ordered by chainl ength (linear alkanes and alkenes) or by chain length and methyl position (mono- and dimethyl alkanes).

We discarded 175 out of 432 cuticular hydrocarbon samples due to contamination and other technical failures, leaving a total of 257 samples from 111 colony replicates that could be used for statistical analysis. Number of used cuticular hydrocarbon samples per colony replicate is available in **Supplementary table 1**.

### Statistical analyses

Prior to running any statistical analysis, we normalized the raw hydrocarbon peak areas of each sample using the log-ratio transformation proposed by Aitchison (1982), as has commonly been used in the analysis of hydrocarbon data:

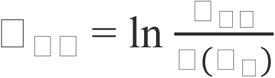

where *y*_*ij*_ is the transformed peak area of the *i*th peak of the *j*th sample, *x*_*ij*_ is the untransformed area of the *i*th peak of the *j*th sample, and *g*(*X*_*j*_) is the geometric mean area of all peaks of the *j*th sample. As in our dataset some samples had zero area values for certain peaks (**Supplementary table 1**), we added a small constant value to each peak prior applying the transformation.

For each colony replicate, we had a single measure for each of the five collective behaviors and for colony productivity but as many as three measures for hydrocarbon peak values. Therefore, we used the mean hydrocarbon replicate value when estimating phenotypic correlations and selection gradients (see below).

We performed all statistical analyses in R version 3.4.1 (R Core Team 2014). A detailed R markdown file including the R scripts, as well as a detailed explanation of each analysis, is included as **Supplementary file 1**.

#### Heritability and genetic correlations estimates of cuticular hydrocarbon compounds

To assess narrow sense heritability (*h*^2^) and genetic correlations (*r*_*G*_) of cuticular hydrocarbon compounds, we analyzed our data using the “animal model” approach (Wilson et al. 2010) with the R package “MCMCglmm” (Hadfield 2010). This mixed-effect model uses a Bayesian Markov chain Monte Carlo (MCMC) approach to decompose phenotypic variance into its genetic and environmental components, allowing the estimation of quantitative genetic parameters. In an animal model, the pedigree of many individuals (i.e. “animals”) is used to make inferences about expected patterns of genetic relatedness among the individuals, and together with observed patterns of phenotypic resemblance among individuals, heritability for measured traits, and genetic correlations between traits are estimated. We treated our replicate colonies (i.e. groups of workers sampled from each replicate colony) as “individuals” in an animal model, and the pedigree of each colony traced the queen and male parents of the worker offspring that made up each colony in the mapping population. Using such an approach, we asked to what degree variation in the expected genetic makeup of workers (i.e. based on the pedigree) predicted the observed phenotypic variation in hydrocarbon profile among groups of workers. While the hydrocarbon profile of each individual worker is expected to depend on both directly on its own genotype (direct genetic effects) and indirectly on the genotypes of its nestmates (through indirect genetic effects) (Linksvayer 2006, van Zweden et al. 2010), our approach does not enable us to estimate the separate contributions of these direct and indirect genetic effects to the observed composite hydrocarbon profile of our replicate worker groups. That is, we cannot quantify the degree to which the hydrocarbon profile of each individual depends on compounds synthesized by that individual, as opposed to compounds synthesized by social partners, but we can quantify the degree to which phenotypic variation in the hydrocarbon profile of groups of workers is predicted by the genotypic makeup of those workers. Similarly, previous animal breeding studies have shown that the total contribution of direct and indirect genetic effects to total genetic variance and total heritability can be estimated by quantifying phenotypic variation among groups of individuals (Peeters et al. 2013; Brinker et al. 2017), although it is not possible to empirically tease apart the separate contribution of direct and indirect genetic effects.

We ran 32 Bayesian univariate models to estimate the heritability of each hydrocarbon compound, and 496 bivariate models (one for each pairwise combination of hydrocarbon variables) to calculate genetic correlations between compounds. Univariate and bivariate models had the same random and fixed effect structure, and differed only in terms of priors specification (**Supplementary file 1**). Since we had multiple hydrocarbon samples for each individual (colony replicate), we ran repeated measures animal models by first partitioning the phenotypic variance into within- and between-individual components. We did so by including an individual identity random term twice in the model: one term (“Colony”) linked individuals to the pedigree structure in order to model the additive genetic variance (*V*_*A*_), whereas the other term (“ID”) was not associated with the pedigree and modelled the “permanent environmental variance” *V*_*PE*_ (Wilson et al. 2010). A third random effect term, named “Block,” was included to account for an extra source of variation due to the fact that samples were collected at different time points from the replicate colonies. To account for the haplodiploid system of our ants, we included in the models the inverse of a sex-chromosomal additive genetic relatedness matrix. This relatedness matrix, which connects colony replicates to their records in the pedigree, reflects the covariance among relatives as a result of the different inheritance patterns of sex chromosomes as compared to the autosomes. This inheritance pattern is the same as what would be expected in haplodiploid organisms. The matrix was built using the function *makeS*() in the package “nadiv” (Wolak 2012). The distribution of the hydrocarbon response variable was set to “gaussian”, and each model ran for 1 000 000 iterations, sampling every 500^th^ data point, and had a burn-in of 10 000 iterations (see **Supplementary file 1**).

Genetic correlation estimates were deemed significant when the 95% confidence intervals of the posterior mode did not span zero.

#### Linear and quadratic selection gradients estimates

We used the two measures of colony productivity (production of sexual and worker pupae) as definition of fitness, and estimated strength and type of selection (e.g., directional, stabilizing, or disruptive) acting on individual hydrocarbons using the regression approach developed by Morrissey and Sakredjda (2013). Briefly, we first estimated the fitness function relating colony productivity to the abundance of a specific hydrocarbon with a generalized additive model (GAM), using the R package “mgcv” (Wood 2017). Then, linear (β) and quadratic (γ) selection gradients are obtained from the fitted GAM model using the function gam.gradients() in the package “gsg” (Morrissey & Sakrejda 2014), which can handle non-normal distributions of fitness. We ran 64 univariate models (32 phenotypic traits × 2 fitness measures). Prior to running the model, hydrocarbon variables were mean-centered and variance standardized. Linear and quadratic selection estimates, case-bootstrapped standard errors and p-values for both definition of fitness are reported in **Supplementary table 2**. Details of model specification can be consulted in **Supplementary file 1**.

#### Correlation between hydrocarbon compounds and collective behaviors

We ran Spearman’s rank-order correlations to evaluate the strength and direction of association between cuticular hydrocarbons and collective behaviors. We ran a model between each compound and each of the five collective behavior (160 models in total). P-values were adjusted using the false discovery rate (FDR) method.

#### Random forest classification analysis

We used a random forest classification analysis (“RF”, Breiman 2001) to determine which cuticular hydrocarbon peaks can better discriminate across the 48 colony genotypes. Although this method does not take into account pedigree relationships, it can provide hints about which hydrocarbons are more variable among colony genotypes, thus highlighting compounds that might be involved in inter- and intra specific recognition. We ran a stratified sampling RF classification model with replacement using the R package “randomForest” (Liaw and Wiener 2001), and we considered hydrocarbon samples from colony replicates belonging to the same colony genotype as part of one of the 48 colony genotypes classes. We used the mean decrease in model accuracy (MDA), as suggested by Cutler (2007), to interpret hydrocarbons importance in classifying the colony genotypes. Model details and specifications can be consulted in **Supplementary file 1**.

## Results

### Heritabilities and genetic correlations

We estimated the heritability of hydrocarbons to be between 0.006 and 0.36, with a median estimated heritability of 0.17 (Figure 1b). We found 31 significant (confidence intervals did not overlap with zero) genetic correlations between hydrocarbons (Figure 2). Although we found many positive and negative genetic correlations, all significant genetic correlations were positive.

**Figure 2.**
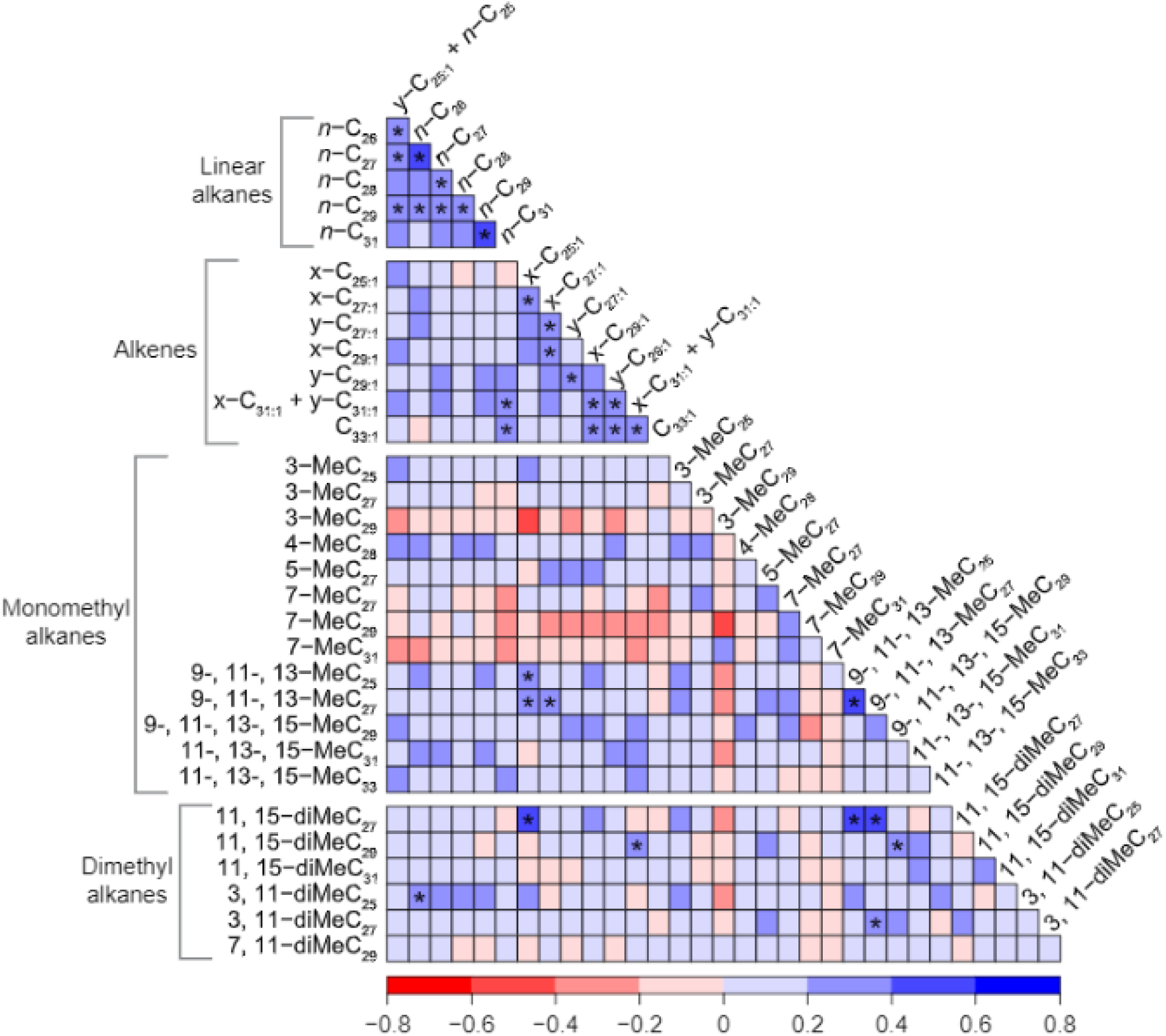
Heatmap showing genetic correlation estimates among cuticular hydrocarbons obtained from bivariate animal models. Compounds are grouped by structural class and ordered by chain length (linear alkanes and alkenes) or by chain length and methyl position (mono- and dimethyl alkanes). Asterisks within cells indicate significant correlations (95% confidence intervals do not span over zero). Different colors indicate the magnitude and direction of the correlation.

### Selection gradients

When defining fitness as the production of reproductives (gynes and males) or workers, we found evidence for significant positive and negative linear selection for many hydrocarbons (Figure 3; see **Supplemental table 2** for estimates and p values). All quadratic selection estimates were not significant.

**Figure 3.**
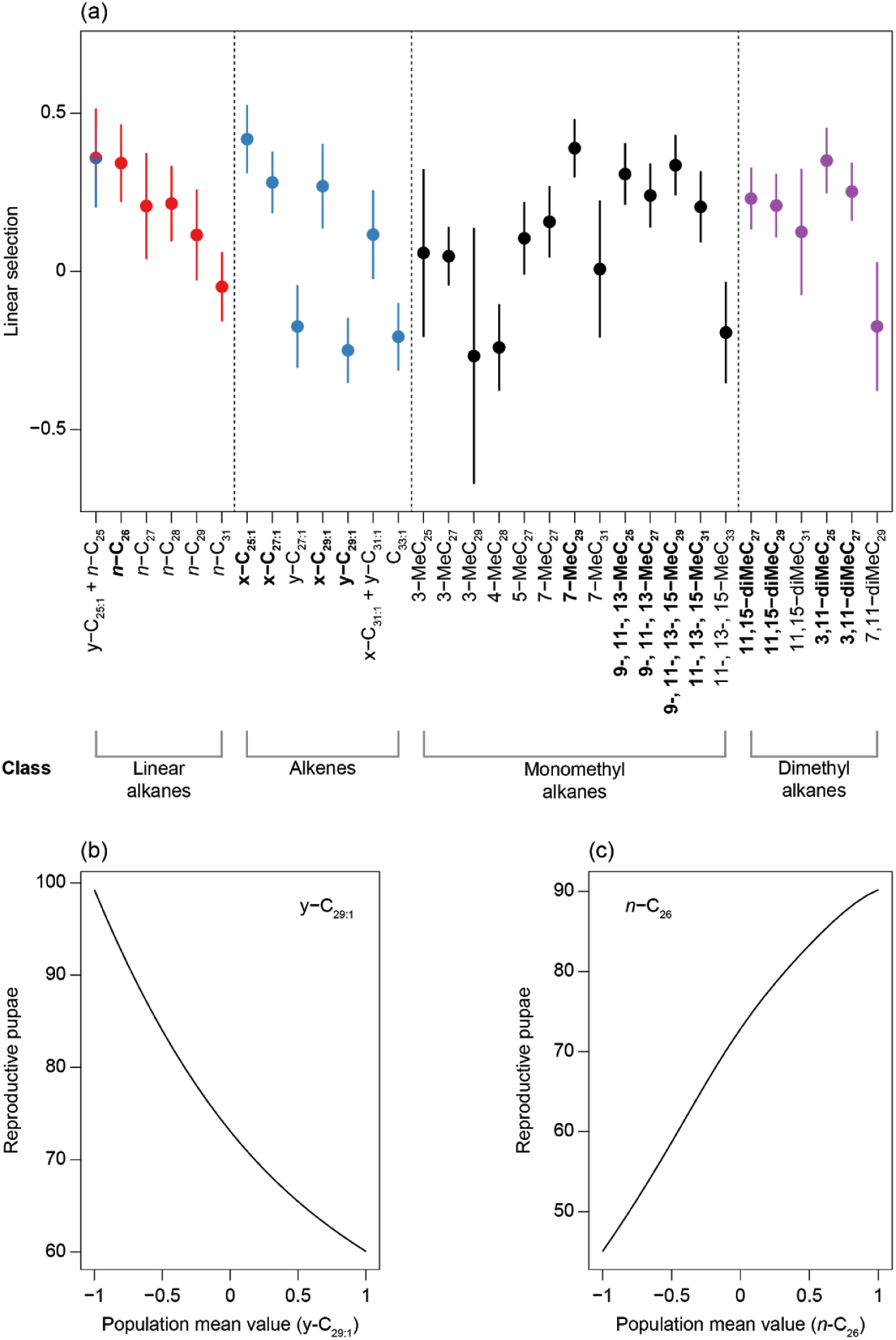
(a) Caterpillar plot showing linear selection estimates, with associated case-bootstrapped standard errors, for individual hydrocarbons using sexual pupae production as measure of colony fitness. Compounds in the plot are grouped and color-coded by structural class and ordered by chain length (linear alkanes and alkenes) or by chain length and methyl position (mono- and dimethyl alkanes). Compounds showing a statistically significant selection gradient are labelled in bold. (b) and (c) show representative fitness landscapes for sexual pupae production as a function of population mean phenotype values of y-C_29:1_ and n-C_26_, respectively.

### Phenotypic correlations

We found many significant phenotypic correlations between individual hydrocarbons and three of the five collective behaviors (foraging, aggression, and group exploration) (Supplementary figure 3).

### Random forest analysis

The accuracy of the RF model was 17.1%, indicating that most of the hydrocarbon samples were correctly assigned to its own colony genotype (**Supplementary file 1**, **Supplementary table 3**). Two compounds, 11,15 diME-C_27_ and 7-MeC_27_, showed the best discrimination accuracy (Supplemental figure 4).

## Discussion

In solitary insects, the heritability of the cuticular hydrocarbon profile is well-characterized (e.g. Ferveur 2005; Thomas & Simmons 2008; Sharma et al. 2012), and patterns of natural and sexual selection acting on cuticular hydrocarbon profiles have also been characterized in several solitary insects (e.g. Blows 2002; Foley et al. 2007; Thomas & Simmons 2008; Berson et al. 2019). In contrast, even though cuticular hydrocarbons are perhaps even more functionally important in social insects -- because they play additional roles in nestmate recognition and in intra-colonial signalling of dominance, reproductive status, and task -- relatively little is known about the heritability of social insect cuticular hydrocarbons (Boomsma et al. 2003; van Zweden et al. 2010) and how they are shaped by natural selection. Here, we begin to elucidate the genetic architecture underlying variation in the hydrocarbon profile and to characterize how selection acts on it in a laboratory population of the ant *Monomorium pharaonis*. We provide evidence that the hydrocarbon profile is heritable, shaped by selection, and that the many hydrocarbons, especially linear alkanes and alkenes, are genetically correlated with each other.

We estimated the total heritability of individual hydrocarbons to be between 0.006 and 0.36, with a median estimated heritability of 0.17 (Figure 1b). These estimates are broadly similar to estimated heritability for collective behavior, body size, caste ratio, and sex ratio made with the same population (Walsh et al. 2019), and are also similar to the range of heritability for individual hydrocarbons estimated from solitary insect populations (e.g. Thomas & Simmons 2008b; Sharma et al. 2012). We expected compounds with high heritability estimates to be among the best at distinguishing between colonies in the random forest analysis because compounds with little genetic component were unlikely to be highly variable between colonies reared in a controlled environment. In accordance with this prediction, the two compounds with the highest heritability estimates (11,15-diMeC27 and 7-MeC27) were also two of the top three compounds at distinguishing between colonies in the random forest analysis. Many of the compounds with relatively high heritability in our study were also highly variable in a previous study of variation in hydrocarbon profiles among 36 *M. pharaonis* colonies collected at sites around the world (Schmidt et al. 2010), although it is difficult to compare our study directly with this previous study because we were able to identify and quantify nearly twice as many hydrocarbon peaks (18 in Schmidt et al. 2010 versus 34 in the current study). Our study, together with this previous study, indicates that heritable variation for many cuticular hydrocarbons is maintained in our mapping population -- which was developed to represent as much genetic and trait variation as possible -- as well as *M. pharaonis* in nature.

As described above, cuticular hydrocarbons play key roles in nestmate recognition, and hence mediate aggression between colonies. Such kin discrimination, changing social behavior depending on genetic relatedness, is widespread from microbes to humans and is central to the evolution of cooperation and inbreeding avoidance (Fletcher et al. 1987; Strassmann et al. 2011; Dudley et al. 2013). However, the degree to which the necessary variation for genetically-based recognition cues is maintained within populations remains broadly unclear. Our results indicate that most compounds making up the cuticular hydrocarbon profile harbor genetic variation that could be informative for genetically-based nestmate recognition or mate choice. Alkenes and monomethyl and dimethyl alkanes are expected to play a larger role in chemical communication (e.g., nestmate recognition) than linear alkanes because they can be distinguished by the position of the double bond or methyl group(s), while linear alkanes can only be distinguished based on their chain length (Gibbs & Pomonis 1995; Akino et al. 2004; Martin et al. 2008). In support of this prediction, previous work in ants found that monomethyl alkanes were more heritable than linear alkanes, suggesting that monomethyl alkanes are better indicators of colony membership (van Zweden et al. 2010). Our results mostly support this prediction as well. In support of the prediction, our random forest analysis revealed that alkenes and monomethyl and dimethyl alkanes had a higher discrimination power than linear alkanes (Supplemental figure 4).

Many pairs of hydrocarbons were positively genetically correlated, and most of these correlations were between two linear alkanes or between two alkenes (Figure 2). Similarly, previous studies in fruit flies found many strong genetic correlations between individual hydrocarbons (e.g. Sharma et al. 2012; Ingelby et al. 2013). It is not surprising that strong genetic correlations exist between many hydrocarbons, especially between compounds of the same structural class, because the production of different hydrocarbons involves many of the same biosynthetic processes (reviewed by Ginzel & Blomquist 2016), and there is expected to be tradeoffs in the production of different structural classes (Foley et al. 2007). These genetic correlations mean that the independent evolution of hydrocarbons will be constrained.

Our study is the first social insect study to link variation in cuticular hydrocarbons with variation in colony productivity, although previous social insect studies have linked variation in cuticular hydrocarbons to worker survival (Sprenger et al. 2018) or to climatic or biotic variation (Menzel et al. 2017; Buellesbach et al. 2018; Sprenger et al. 2018). Because *M. pharaonis* queens cannot form new colonies without workers (i.e. colonies reproduce by budding; Passera 1994; Buczkowski & Bennett 2009), we defined fitness in two ways: the number of new reproductives (gynes and males) produced or the number of new workers produced. Interestingly, we found similar linear selection patterns using both definitions, as all significant linear estimates were in the same direction between the two definitions (Figure 3, **Supplement table 2**). This suggests that the hydrocarbon profile optima are largely aligned for the production of both reproductives and workers in our study population.

These selection results beg the question: what is the likely causal link between variation in worker cuticular hydrocarbons and variation in colony productivity in our study population? Interestingly, cuticular hydrocarbon profile was phenotypically correlated with collective behavior (Supplemental figure 3), in particular foraging rate, which in turn was also positively associated with colony productivity (Walsh et al. 2019). This relationship between variation in cuticular hydrocarbon profile, foraging rate, and colony productivity could be mediated by effects of hydrocarbons on the desiccation resistance of workers. However, our colonies likely experienced relatively low water stress since the colonies were kept in climate controlled chambers at 50% humidity, and the colonies always had access to water. Alternatively, worker hydrocarbon profiles might influence colony-level division of labor or task allocation, which could in turn influence foraging rate and colony productivity. As described above, in addition to effects on desiccation resistance, ant cuticular hydrocarbons are well known to influence nestmate recognition and inter-colonial aggression in many ant species, including *M. pharaonis* (Pontieri 2014). However, colonies in our study were isolated from each other throughout the course of the study, so that differences in the outcome of aggressive encounters between colonies that likely contribute to differences in nature for colony survival and productivity (Reeve and Hōlldobler 2007) cannot explain the patterns of selection on hydrocarbon profile that we observed in our laboratory population. Similarly, while cuticular hydrocarbon profiles may also mediate mate choice in *M. pharaonis*, such a mechanism could not explain the association between worker hydrocarbons and colony productivity that we observed.

An interesting complication of the genetic architecture of social insect cuticular hydrocarbon profiles (Linksvayer 2006, 2015) is that the social environment experienced by each individual within a social insect colony strongly influences its hydrocarbon profile, since hydrocarbons are mixed throughout the colony via trophallaxis and allogrooming between colony members (Bonavita-Cougourdan et al. 1997; van Zweden et al 2009, 2010; Martin et al. 2012; Khidr et al. 2013; Gevar et al. 2017). As a result, the genetic architecture of the hydrocarbon profile, like other socially-influenced traits, depends on the collective genetic makeup of colonies (Linksvayer 2006, 2015). That is, the hydrocarbon profile of each individual worker depends directly on its own genotype (direct genetic effects; e.g., via hydrocarbons it synthesized) but also depends indirectly on the genotypes of its nestmates (indirect genetic effects; e.g., via hydrocarbons synthesized by nestmates and transferred to the focal individual’s cuticle via social trophallaxis, grooming, etc.). Because we quantified the cuticular hydrocarbon profile of *groups* of workers from each colony, we were not able to distinguish between hydrocarbons that were readily transferred among nestmates and those that were only produced by a subset of workers and not transferred (see van Zweden et al. 2010), or to separately estimate the contribution of variation in direct and indirect genetic effects to estimated total heritability (Moore et al. 1997; McGlothlin et al. 2010; Linksvayer 2015). Furthermore, we were not able to consider differences in cuticular hydrocarbons between individual workers based on age, task within the colony, or differences in genotypes within a colony.

We conducted the current study in a laboratory environment, which enabled us to strictly control the colony demography (i.e. queen number, worker number, etc.), diet, and environmental conditions experienced by the colonies. Such control in particular is valuable given the complexity of social insect colonies (Linksvayer 2006; Kronauer & Libbrecht 2018) and the sensitivity of the hydrocarbon profile to changes in the environment or diet (Francis et al. 1989; vander Meer et al. 1989; Liang & Silverman 2000; Tissot et al. 2001; Buczkowski et al. 2005; van Zweden et al. 2009; Pavković-Lučić et al. 2016; Menzel et al. 2017; Buellesbach et al. 2018). Although future field studies would be beneficial, in particular to identify how variation in cuticular hydrocarbons affects colony productivity in a natural setting, a field study on a similar scale as our study is likely not feasible.

Overall, this study increases our understanding of the genetic architecture of the hydrocarbon profile and demonstrates that the hydrocarbon profile is shaped by natural selection. Although numerous genes underlying variation in the hydrocarbon profile have been identified in *Drosophila* (Dembeck et al. 2015), the hydrocarbon profile performs different functions in the social insects and therefore future studies should focus on determining whether the same genes are involved in the expression of social insect hydrocarbon profiles, and how variation in these genes affects variation in the social insect hydrocarbon profile. For example, a recent study used a candidate gene approach and found that inotocin, a peptide similar to oxytocin/vasopressin, regulates the production of hydrocarbons in the ant *Camponotus fellah* (Koto et al. 2019). Future studies should use unbiased approaches such as quantitative trait locus (QTL) mapping in mapping populations (e.g. Valdar et al. 2006) such as ours, and association mapping in natural populations (e.g Kocher et al. 2018).

---

**Supplementary figure 1.**
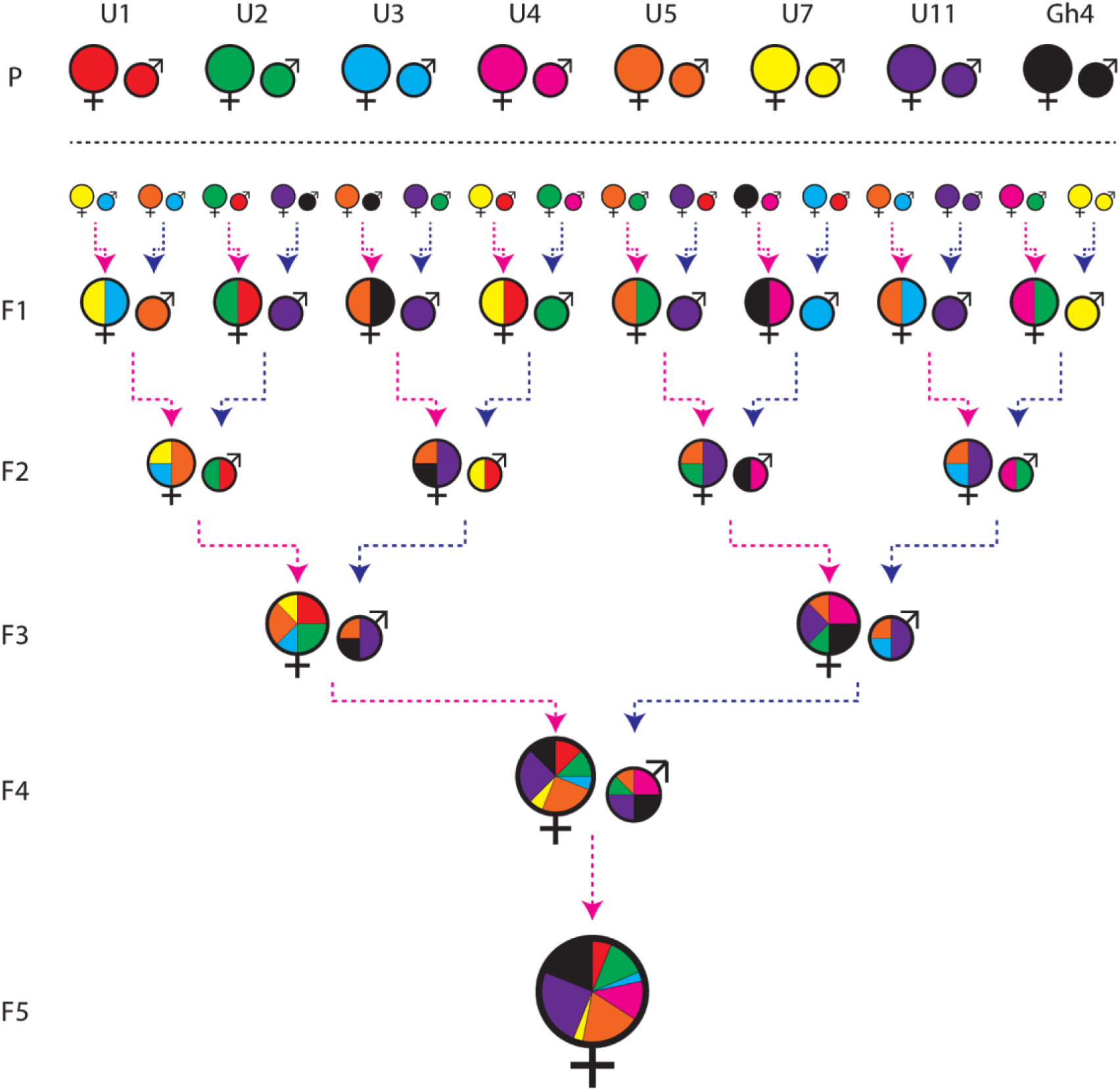
Crossing scheme employed for the creation of the *M. pharaonis* mapping population. For simplicity, only the crosses needed to create a generation F5 colony are showed, although eight parental colonies (P) were sequentially intercrossed for nine generations. The black dashed horizontal line separates generations of inbreeding and the onset of the crossing procedure. Briefly, gynes (♀) and males (♂) from each of the eight parental colonies (colour coded) were collected and crossed with sexuals from other parental colonies. From the resulting F1 offspring, new gynes and males were collected (pink and blue dashed arrows, respectively) and crossed in order to give birth to the F2 generation. The same protocol was applied for the subsequent generations. The expected genetic contribution of each parental colony to the genotype of each individual represented in the figure is indicated by different sizes of pie chart slices, colour coded according to parental colony.

**Supplementary figure 2.**
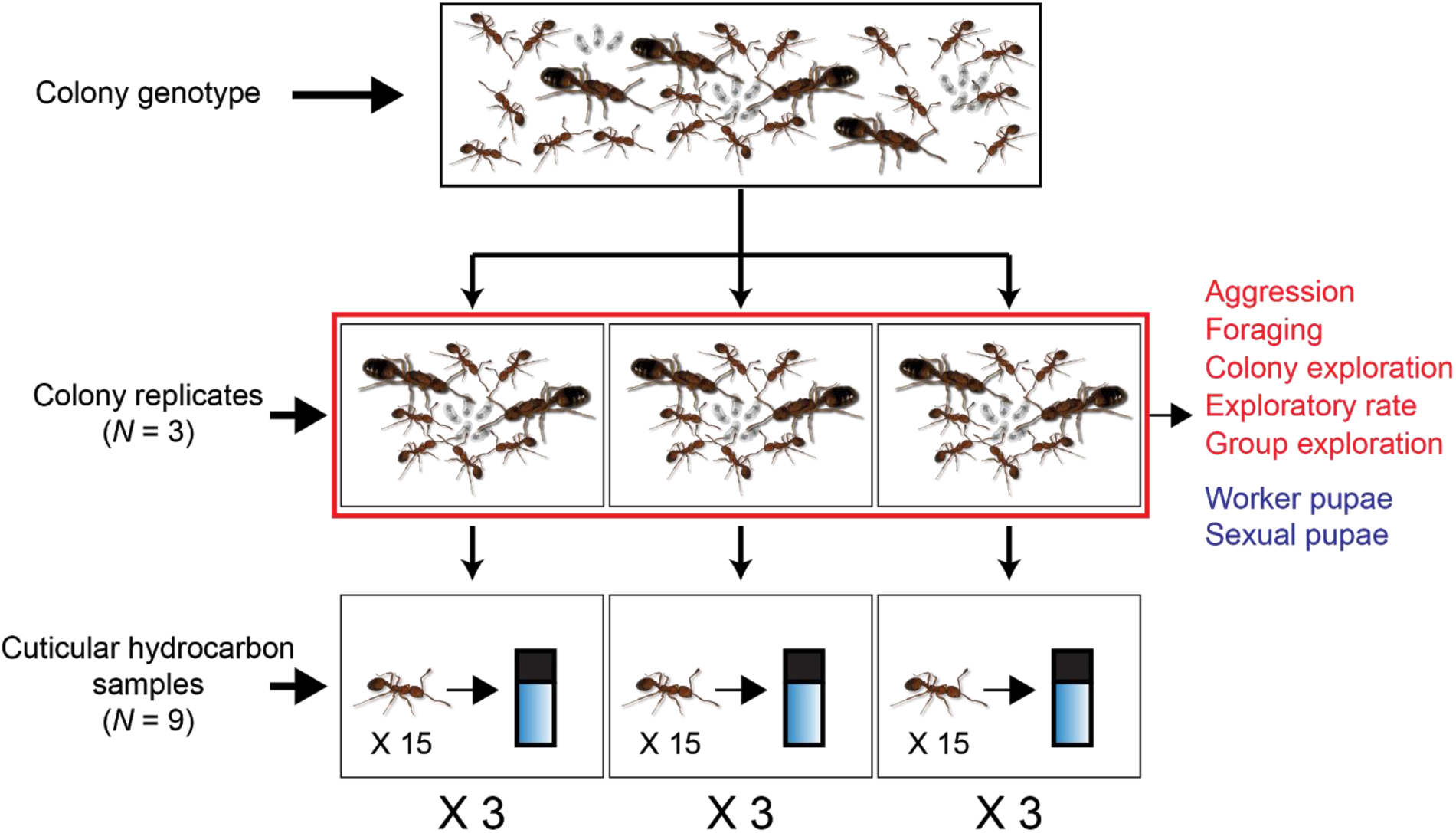
Experimental design and sampling scheme for one of the 48 colony genotypes. Note that the number of queens, workers and brood shown is not meant to represent the actual composition of the colonies (see the methods section for information about colony replicates demographics).

**Supplementary figure 3.**
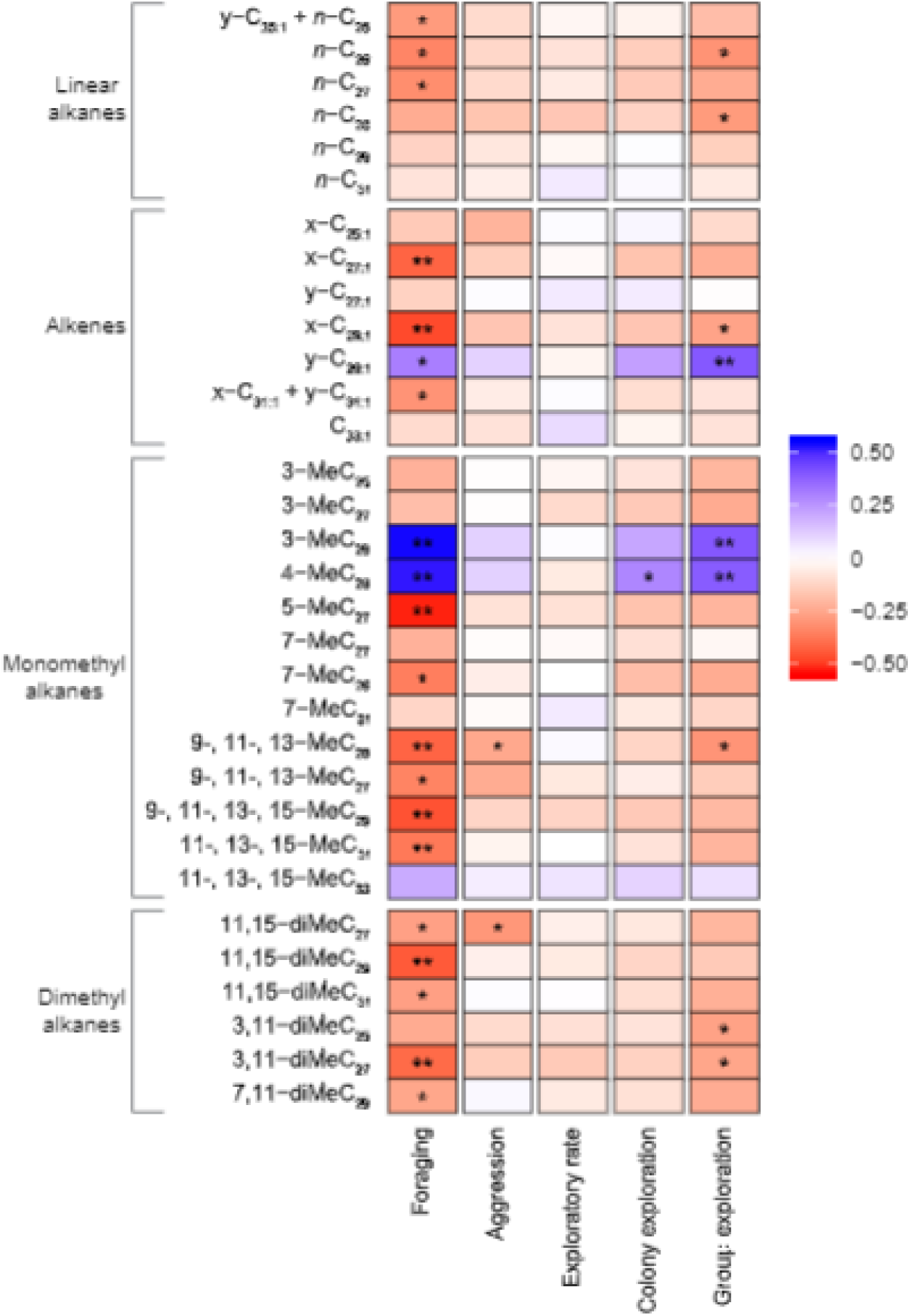
Heatmap showing phenotypic correlations between the five collective behaviors and individual hydrocarbons. Compounds are grouped by structural class and ordered by chain length (linear alkanes and alkenes) or by chain length and methyl position (mono- and dimethyl alkanes). Cell color represents magnitude and sign of the Spearman rank correlation coefficient. Asterisks within cells represent statistically significant associations (“*” = p < 0.05; “**” = p < 0.001; false discovery rate adjusted p-values).

**Supplementary figure 4.**
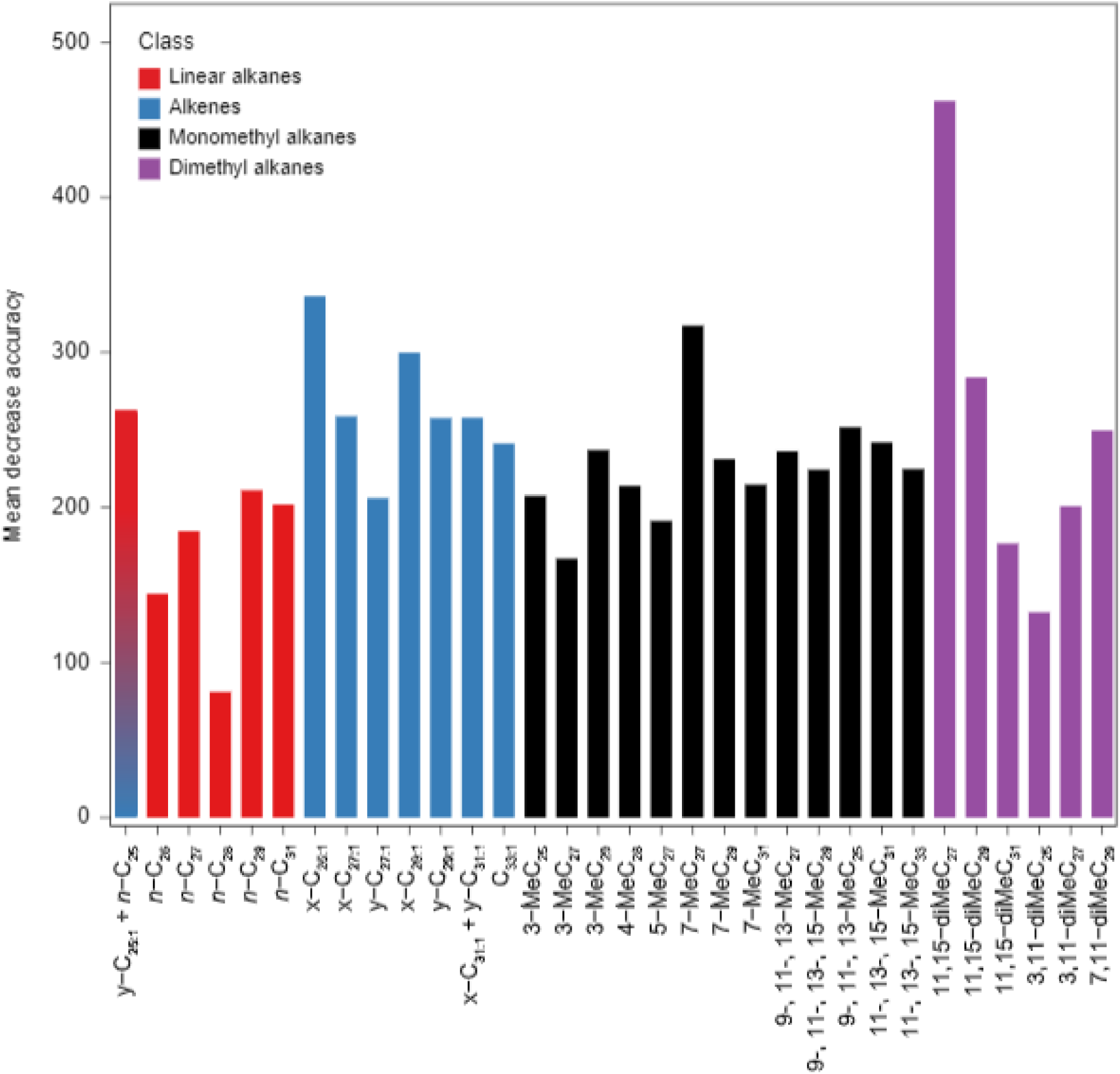
Random forest variable importance plot showing the power of individual compounds to discriminate among colony genotypes (expressed as mean decrease in model classification accuracy when the variable is not included). Higher values indicates higher discrimination power. Compounds in the plot are grouped and color-coded by structural class and ordered by chain length (linear alkanes and alkenes) or by chain length and methyl position (mono- and dimethyl alkanes).

**Supplementary table 1.** Data set

**Supplementary table 2.** Linear and quadratic selection estimates for individual hydrocarbons using either sexual or worker pupae production as colony fitness measure.

**Supplemental table 3**. Confusion matrix generated by the random forest analysis. Cells highlighted in green represent the number of hydrocarbon samples correctly assigned to its own colony genotype. Cells highlighted in red indicate the number of incorrect assignments, as well as to which colony genotype they were incorrectly assigned. Adjacent to the matrix, we report the number of hydrocarbon samples for each genotype (# samples); the number of samples randomly drawn from each colony genotype to build each tree in the RF analysis (#sample/tree); the number of the Out-Of-Bag samples for each colony genotype (OOB samples); the number of samples correctly assigned to its own colony genotype (Correct); the number of incorrect assignments (Incorrect); the Out-Of-Bag assignment error rate for each colony genotype (Class Error).

**Supplementary file 1.** R markdown pdf file containing the R scripts used for the statistical analyses, as well as step-by-step description and in-depth explanation of each line code. The file includes also model diagnostic scripts.

