## Supplemental File 1 for "Ant cuticular hydrocarbons are heritable and associated with variation in colony productivity"

### R code for the analyses performed in Walsh et al. 2019

#### 1) Heritability of CHC compounds

##### Load packages

```
library(MCMCglmm)
library(nadiv)
```

##### Load datasets

```
PEDIGREE=read.table("Genotype Pedigree - Heritability.txt", header=T)# genotype pedigree
PEDIGREE.COLONY=read.table("Colony Pedigree - Heritability.txt",header=T)# colony info
CHC=read.table("CHC - Heritability.txt",header=T)# log-ratio transformed CHCs areas
```

##### Encode CHC dataset variables

create the "animal" variable (MCMCglmm will recognize the term animal and use it to relate individuals to their records in a pedigree); create "ID" variable (same as "animal" variable) to model permanent environment effects in an animal model to prevent upward bias in the additive genetic variance (Va).
"Animal" and "ID" factors partition additive and permanent environment effects.
N.B. "ID" variable is required for models including repeated measures!

```
CHC$Block<-as.factor(CHC$Block)# convert block vector into factor
CHC$Colony<-as.factor(CHC$Colony)# convert colony vector into factor
CHC$animal<-interaction(CHC$Genotype,CHC$Colony,drop=T)# "animal" variable
CHC$ID<-CHC$animal# "ID" variable
```

##### Encode colony info dataset variables

```
PEDIGREE.COLONY$Block<-as.factor(PEDIGREE.COLONY$Block)# convert block into factor
PEDIGREE.COLONY$Colony<-as.factor(PEDIGREE.COLONY$Colony)# convert colony into factor
```

##### Create the final pedigree dataset

"animal" factor is the same as the "animal" factor in the CHC dataset.

```
NEW.PEDIGREE=merge(PEDIGREE.COLONY,PEDIGREE,by.x="Genotype",by.y="animal",all.X=TRUE)
NEW.PEDIGREE$animal=interaction(NEW.PEDIGREE$Genotype,NEW.PEDIGREE$Colony,drop = T)
NEW.PEDIGREE=NEW.PEDIGREE[c("animal","dam","sire","sex")]
PEDIGREE<-PEDIGREE[c("animal","dam","sire","sex")]
FINAL.PEDIGREE=rbind(NEW.PEDIGREE,PEDIGREE) # pedigree
rm(NEW.PEDIGREE,PEDIGREE,PEDIGREE.COLONY)
```

##### Create the inverse of the additive genetic relationship matrix

Haplodiploidy is modelled here in the same way as a sex chromosome, with male being the heterogametic sex.

```
Mat_A <- makeS(prepPed(FINAL.PEDIGREE),heterogametic="M",returnS=T)
```

##### Formulate univariate priors for the model

R for the residual variance and G for the random effects.
Often used priors correspond to an inverse-Gamma distribution with shape and scale parameters equal to 0.01. However, this inverse-Gamma could become unwantedly 'informative' when variance components

are close to 0 (see Gelman 2006), which was our case for certain CHC compounds. We ran models with different prior specifications (not showed),until we found priors that ensured convergence of the model and no autocorrelation. The final priors are reported below.

```
prior <- list(R = list(V=1, nu=1.002),
              G = list(G1 = list(V=1, nu=1.002),
              G2 = list(V=1, nu=1.002),
              G3 = list(V=1, nu=1.002)))
```

##### The MCMCglmm model

We ran a univariate model for each CHC. The response variable was modelled as a normal distributed variable. We included Colony (the "animal" factor), Block and ID as random effects.Burning step of 10000; thinning of 500; 1 million iterations.We provide an example for peak 1.

```
MCMC <- MCMCglmm(Peak1~ 1, random=~ ID+animal+Block,
                 ginverse=list(animal=Mat_A$Sinv),
                 nitt = 1e6, burnin = 1e4, thin = 5e2,
                 family = c(rep("gaussian",1)),
                 pl = T , pr = T, data = CHC,prior = prior)
```

##### Model diagnostic

**1.** We first checked that the autocorrelation across montecarlo chains for all variables was as close to zero as possible (values below 0.1 are considered acceptable) for all lag values greater than zero.We used the function autocorr()

```
autocorr(MCMC$VCV)
```

```
## , , ID
##
##                     ID       animal         Block        units
## Lag 0      1.000000000 -0.157103444 -0.0005260406 -0.092300542
## Lag 500   -0.005099924 -0.003883295 -0.0085782112 -0.065576245
## Lag 2500  -0.029179136  0.028762015 -0.0191532483 -0.013488620
## Lag 5000   0.034648431 -0.052874734  0.0105464052 -0.002717903
## Lag 25000 -0.005854540 -0.016907360 -0.0064787536  0.033367000
##
## , , animal
##
##                     ID      animal       Block        units
## Lag 0     -0.157103444  1.00000000  0.04079926 -0.032043043
## Lag 500    0.074597245 -0.04461340 -0.02309834  0.034836911
## Lag 2500   0.018382267 -0.03748974  0.01167241  0.012009439
## Lag 5000  -0.019083873  0.02996178 -0.02082077 -0.002021049
## Lag 25000  0.002332826  0.02142337  0.04489260 -0.008085888
##
## , , Block
##
##                      ID       animal         Block        units
## Lag 0     -0.0005260406  0.040799256  1.0000000000 -0.011909949
## Lag 500   -0.0203060904 -0.014594113 -0.0185749199 -0.012532803
## Lag 2500  -0.0473797350  0.022602595 -0.0028578755 -0.006540692
## Lag 5000  -0.0007209164 -0.014499845  0.0483786634  0.019498932
## Lag 25000  0.0204830436  0.003082727 -0.0008981349  0.021175503
##
```

```
## , , units
##
##                         ID      animal        Block        units
## Lag 0     -0.0923005420 -0.03204304 -0.011909949  1.000000000
## Lag 500   -0.0209359401 -0.01263845  0.000735971  0.014270281
## Lag 2500   0.0005233007 -0.03214365 -0.047601789  0.064046776
## Lag 5000   0.0338658693 -0.01957439 -0.024443324  0.009429890
## Lag 25000 -0.0062129175 -0.03822576 -0.001887609 -0.007240275
```

**2.** We checked for convergence using the function heidel.diag()

```r
heidel.diag(MCMC$VCV)
```

```
##
##        Stationarity start     p-value
##        test         iteration
## ID     passed       1         0.670
## animal passed       1         0.925
## Block  passed       1         0.742
## units  passed       1         0.478
##
##        Halfwidth Mean   Halfwidth
##        test
## ID     passed    0.0737 0.000746
## animal passed    0.1090 0.001257
## Block  passed    0.2898 0.007177
## units  passed    0.0610 0.000310
```

**3.** We visually inspected the well mixing of chains for the "animal" variable using the function traceplot(). If chains mixed well, the graph should look as a fuzzy caterpillar, without any visible trend.

```r
traceplot((MCMC$VCV[, "animal"])/(MCMC$VCV[, "ID"] +
           MCMC$VCV[, "units"]+MCMC$VCV[, "animal"] +
           MCMC$VCV[, "Block"]))
```

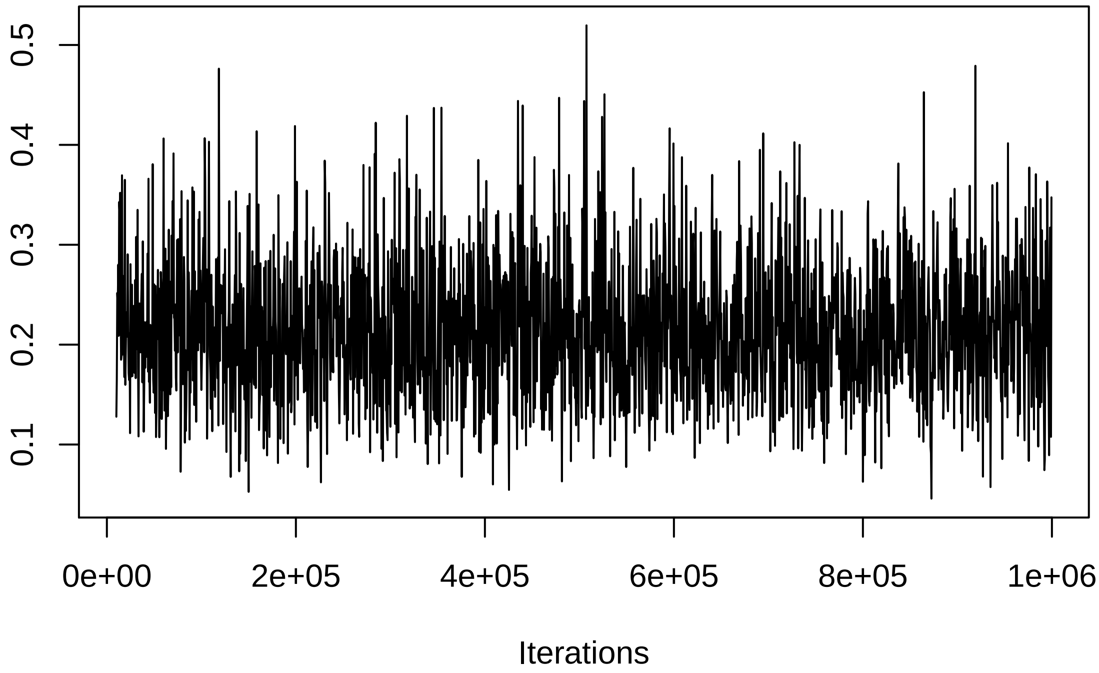


**4.** We visually inspected the posterior density plot of the "animal" parameter using the function densplot(). The distribution, ideally, should look unimodal.

```
densplot((MCMC$VCV[, "animal"])/(MCMC$VCV[, "ID"] +
          MCMC$VCV[, "units"]+MCMC$VCV[, "animal"] +
          MCMC$VCV[, "Block"]))
```

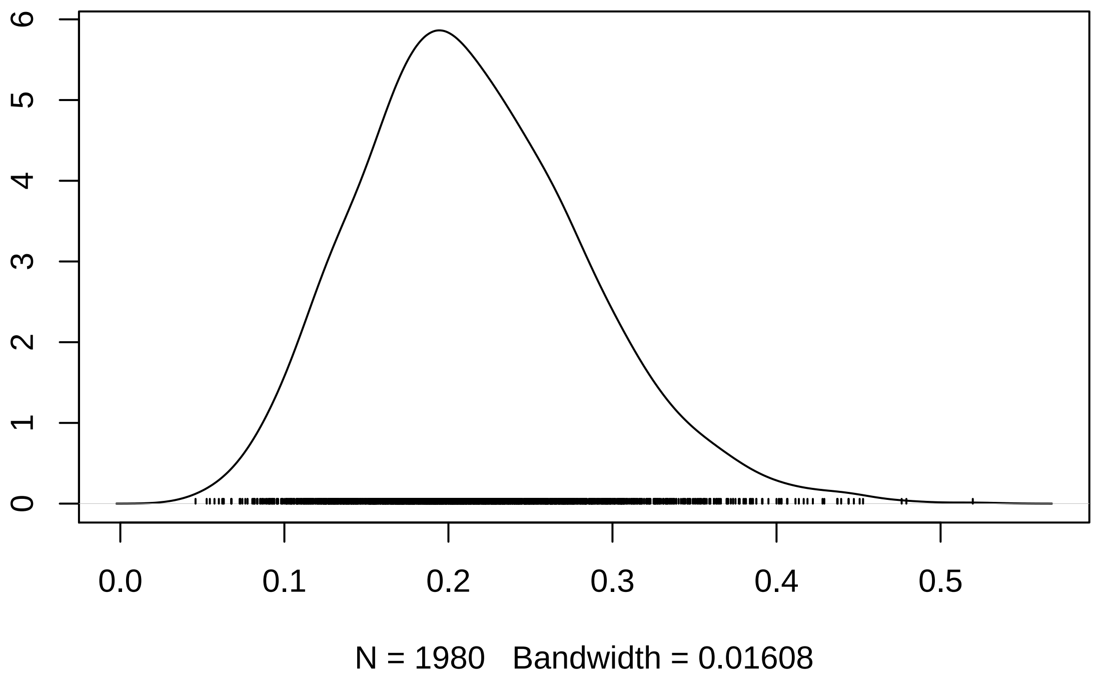


**5.** We ensured that the model retained enough independent runs (at least 1500) by looking at the effective sample size for the "animal" parameter.The maximum number of independent runs that our model can retain in absence of autocorrelation is 1980.

```
effectiveSize((MCMC$VCV[, "animal"])/(MCMC$VCV[, "ID"] +
              MCMC$VCV[, "units"]+MCMC$VCV[, "animal"] +
              MCMC$VCV[, "Block"]))
```

```
## var1
## 1980
```

**Calculate posterior heritability and the associated 95% confidence intervals**

**Posterior heritability**

```
posterior.mode((MCMC$VCV[, "animal"])/(MCMC$VCV[, "ID"] +
               MCMC$VCV[, "units"]+MCMC$VCV[, "animal"] +
               MCMC$VCV[, "Block"]))
```

```
##      var1
## 0.1926672
```

**Confidence intervals**

```
HPDinterval((MCMC$VCV[, "animal"])/(MCMC$VCV[, "ID"] +
            MCMC$VCV[, "units"]+MCMC$VCV[, "animal"] +
            MCMC$VCV[, "Block"]))
```

```
##            lower     upper
```

```
## var1 0.08111108 0.3474819
## attr(,"Probability")
## [1] 0.95
```

For each CHC, we repated the code outlined above by simply replacing the peak number in the model.
